# Individual differences in Game Transfer Phenomena are associated with creativity and positive schizotypy but not with perceptual priming

**DOI:** 10.64898/2026.10.27.762403

**Authors:** Julio C. Llamas-Alonso, Lia Kvavilashvili, George Georgiou, Angelica B. Ortiz de Gortari

## Abstract

Game Transfer Phenomena (GTP) refer to involuntary experience of video-game elements as part of gamers’ everyday perception, cognition, and behaviour. Although previous research has identified Internet Gaming Disorder (IGD), positive schizotypy, and spontaneous cognitions (e.g., mind-popping, daydreaming) as key predictors of GTP, the role of implicit memory processes and creativity remains unclear. The present study examined whether these cognitive processes are associated with GTP frequency while replicating established predictors such as IGD, involuntary cognitions, and schizotypal traits. A total of 85 video game players (53% males; *M*_age_ = 24.89, *SD* = 5.87) completed a word fragment completion task to assess individual differences in perceptual implicit memory (priming). To measure creativity, participants completed the Alternate Uses Task (AUT) as a measure of divergent thinking, and a free association task to estimate forward flow, a robust predictor of creative cognition. Results showed robust priming effects at the group level (*M* = .23, *SD* = .14), however, individual differences in perceptual priming were not associated with GTP frequency. In contrast, AUT scores were positively associated with GTP and contributed uniquely to its prediction. A hierarchical regression model explained 58.2% of the variance in GTP frequency, with IGD, daydreaming, and positive schizotypy as the strongest predictors, alongside hours played per session and AUT scores. These findings suggest that GTP may not be driven by domain-general perceptual implicit memory processes but may be more strongly linked to individual differences in spontaneous cognitions and conceptual associative priming processes.

## Introduction

Game Transfer Phenomena (GTP) refer to a range of involuntary experiences in which elements from video games transfer into real-life perception, cognition, and behaviour [1]. These experiences can include inner and outer visual and auditory intrusions, altered perceptions, automatic thoughts, and behavioral tendencies influenced by recently played games [2]. For instance, a player may involuntarily hear a game’s repetitive soundtrack hours after turning off the console, momentarily perceive real-world architecture as interactive, climbable obstacles typical of open-world environments, or execute an automatic physical movement learned during the gameplay [3–5]. Appendix provides a brief overview of common examples documented in the literature on GTP.

Due to their transient and non-pathological nature, GTP have increasingly been recognized as being part of a broader domain of spontaneous cognitive experiences [6,7]. From a cognitive perspective, GTP share multiple characteristics with other involuntary mental phenomena such as mind-popping, mind-wandering, and involuntary autobiographical memories. These experiences are typically spontaneous, non-deliberate, occur during relatively undemanding activities, and are often triggered by internal or external cues [8–10]. Previous research has shown that such experiences are associated with individual differences in cognitive processing, including daydreaming, fantasy proneness, attentional control, and susceptibility to intrusive thoughts [11–16].

Building on this perspective, researchers have increasingly examined why some individuals are more susceptible to experiencing GTP than others. Evidence accumulated over the past few years suggests that GTP are associated with both cognitive and gaming-related variables, although the mechanisms underlying these associations remain poorly understood. One theoretical account proposed to explain the occurrence of GTP is the long-term priming hypothesis [7]. Priming refers to the facilitation of processing or retrieval following prior exposure to stimuli, often occurring without conscious awareness of the influence of the prior experience [17–19]. Within the context of involuntary autobiographical memories and mind-popping, long-term priming has been proposed as an important mechanism that determines the likelihood with which a particular memory or mind-pop can emerge into consciousness [8,20,21]. These studies have demonstrated that recently encountered stimuli can increase the likelihood of subsequent involuntary recall, even in the absence of intentional retrieval [21].

In line with this framework, recent diary-based research on GTP demonstrated that most GTP experiences occurred shortly after gameplay exposure, with participants frequently reporting recent contact with the game related to the GTP experience [7]. Moreover, GTP experiences were commonly triggered by identifiable internal or external cues and tended to emerge during cognitively undemanding activities, conditions that closely resemble those observed in studies of involuntary autobiographical memories and mind-popping [8,21]. These findings provide ecologically valid support for long-term priming accounts of involuntary cognition by suggesting that recent gameplay exposure may temporarily increase the accessibility of gaming-related semantic and autobiographical representations. However, despite this indirect evidence, no study to date has directly examined whether individual differences in priming ability are associated with the frequency of GTP experiences.

Recent research on autobiographical memory priming further suggests that spontaneous cognitions may emerge through associative activation within autobiographical memory networks [22]. According to semantic-to-autobiographical memory priming model [23,24], everyday semantic processing continuously activates autobiographical memory representations through spreading activation mechanisms [25]. Studies on reminiscence priming have demonstrated that deliberately recalling autobiographical events from a particular time-period can increase the subsequent occurrence of involuntary memories that are associatively related, but not identical, to previously activated memories [20,21]. Similarly, recent studies have shown that autobiographical memories can be primed through a broad range of semantic and contextual stimuli, including words, images, sounds, and narratives [26,27]. For example, Mace and Aaron [27] demonstrated that narrative processing itself can prime involuntary autobiographical memories, extending semantic-to-autobiographical priming beyond isolated lexical or perceptual cues. Participants who read short narratives subsequently reported more involuntary autobiographical memories during a vigilance task compared to control participants. Importantly, these findings suggest that autobiographical memory activation may emerge from naturalistic and meaningful cognitive processing contexts rather than from simple perceptual repetition alone.

This distinction may be particularly relevant for GTP, as many GTP experiences involve spontaneous reinterpretations, associations, and context-dependent activations of gaming-related representations rather than direct repetitions of recently encountered stimuli. From this perspective, GTP may reflect spreading activation within gaming-related autobiographical and semantic networks triggered by recent gameplay exposure. This possibility is also consistent with evidence showing that repeated naturalistic exposure to contextually relevant stimuli can produce persistent increases in conceptual accessibility outside laboratory settings [28], suggesting that prolonged engagement with gaming environments may maintain game-related representations in a heightened state of activation. Video games may constitute particularly potent semantic-to-autobiographical priming environments because they involve repeated multisensory exposure, emotional salience, immersive narrative engagement, and highly interactive cognitive processing.

In addition to implicit memory processes, creativity may also play a role in the frequency of GTP. Creativity is commonly defined as the ability to generate novel and useful ideas and has been linked to flexible associative processing and access to remote semantic networks [29]. Individuals high in creativity tend to produce more diverse and unconventional associations, which may increase the likelihood of unusual cognitive experiences. Previous research has also reported associations between creativity and spontaneous cognition, including mind-popping and divergent thinking [30]. Consequently, measures of divergent thinking provide a useful framework for investigating whether individual differences in creative associative processing contribute to the occurrence of GTP.

One of the most widely used measures of divergent thinking is the Alternate Uses Task (AUT) [31], in which participants generate as many original uses as possible for common everyday objects within a limited time period. Performance on the AUT has shown consistent associations with multiple measures of creativity, for a meta-analytic review see [29], making it one of the most widely accepted experimental measures of creative cognition. Traditional AUT scoring typically relies on indices such as fluency of output, flexibility, and originality or uniqueness of responses within a sample that is being tested [32]. However, these approaches have been criticized because uniqueness scores are influenced by both response fluency and sample size, making it difficult to distinguish genuinely creative ideas from statistically infrequent responses [33]. Moreover, statistical rarity does not necessarily imply creativity, as uncommon responses may lack appropriateness or practical value. To address these limitations, George and Wiley [34] modified the AUT by instructing participants to avoid conventional uses of the target object and instead generate only highly original alternatives. By discouraging readily accessible responses, this version encourages deeper semantic search and more flexible associative processing, potentially providing a more sensitive measure of creative idea generation than traditional AUT instructions.

Another relevant approach as a measure of creativity is the forward flow coefficient derived from free association tasks, which quantifies the semantic distance between successive thoughts within a stream of associations [35]. Unlike traditional divergent thinking measures, forward flow captures the dynamic progression of thought by estimating how far each new idea departs semantically from preceding ideas within a semantic network. Gray et al. [35] demonstrated that higher forward flow scores reflected greater semantic divergence and more flexible associative processing, whereas lower scores indicated more constrained and semantically similar thought patterns. Also, participants with higher coefficients of forward flow obtained higher scores in multiple creativity tasks [35]. Given that GTP involve the spontaneous activation and recombination of game-related representations in real-world contexts, forward flow may provide a sensitive index of the associative mechanisms that facilitate these experiences. Therefore, examining forward flow alongside traditional measures of creativity may help clarify the role of dynamic associative processes in the emergence of GTP.

Closely related to creativity are schizotypal traits, particularly positive schizotypy, which involve unusual perceptual experiences, magical thinking, and cognitive-perceptual distortions [36]. A growing body of research suggests that creativity and schizotypal traits may share underlying cognitive mechanisms, including reduced latent inhibition and increased associative processing. Positive schizotypy has also been consistently linked to GTP in recent studies [6,7], with individuals higher in these traits reporting more frequent GTP experiences.

Several other individual-difference variables have also been found to predict GTP frequency. For example, Internet Gaming Disorder (IGD) has consistently been identified as one of the strongest behavioral predictors, likely reflecting greater exposure to game-related stimuli and deeper cognitive immersion. Spontaneous cognitive tendencies such as daydreaming and mind-popping have also shown reliable associations with GTP, further supporting the view that GTP form part of a broader class of involuntary cognitive experiences [6,7].

Despite these advances, several important gaps remain. First, the role of implicit memory processes such as priming has not been directly examined in relation to GTP. Second, although creativity and divergent thinking have theoretically been linked to spontaneous cognition and schizotypy, their relationship with GTP remains largely unexplored. Third, it remains unclear how these cognitive variables interact with established predictors such as IGD and positive schizotypy in explaining individual differences in GTP.

Therefore, the present study aimed to examine the relationships between priming, creativity, and GTP while simultaneously replicating and extending previous findings on established predictors such as IGD, spontaneous cognition, and positive schizotypy [6,7]. Specifically, the study investigated whether individual differences in priming performance are associated with GTP frequency, whether creativity contributes to the prediction of GTP, and whether these effects persist after controlling for gaming behaviour, spontaneous cognitive processes, and schizotypal traits.

To address these aims, participants completed several questionnaires measuring GTP, as well as spontaneous cognition, IGD, and schizotypal traits. They also completed creativity tasks (AUT and a free association task) and a standard implicit memory paradigm consisting of a study phase, a filler interval, and a subsequent test phase was employed [37]. Implicit priming was assessed using a word-fragment completion task, in which participants completed partially presented words without being informed that some fragments corresponded to words encountered during the study phase. Priming scores were calculated as the difference between completion rates for studied and non-studied items [38]. The word-fragment completion task was selected because it is among the most widely used measures of perceptual priming and has been shown to produce robust priming effects over relatively long retention intervals while minimizing contamination from explicit recollection [39]. This paradigm allowed us to examine whether individual differences in GTP frequency were associated with variations in implicit memory performance.

**Hypothesis 1** In line with previous literature, suggesting that priming influences spontaneous cognitive experiences, it was expected that priming scores would be positively associated with the frequency of GTP.

**Hypothesis 2** Given that creativity has been linked to spontaneous and automatic cognitive processes, it was predicted that creativity would be positively associated with GTP frequency and would contribute uniquely to its prediction beyond other related variables.

**Hypothesis 3** If GTP shares underlying mechanisms with other spontaneous cognitive phenomena and is associated with psychological traits, then Internet Gaming Disorder (IGD), spontaneous cognitive processes (i.e., mind-popping and daydreaming), and positive schizotypy would emerge as significant predictors of GTP frequency in a hierarchical regression model.

## Method

### Participants

A power analysis was conducted using G*Power [40] to determine the required sample size for detecting a medium effect size (*f²* = .15) with a statistical power of .90 and 12 predictors. The analysis indicated a minimum sample of 88 participants. To account for potential exclusions, 120 participants were initially recruited. To take part, participants were required to meet the following inclusion criteria. First, all participants had to be active video game players, defined as engaging in any type of video game for at least 4 hours per week on a regular basis over the last 12 months. Second, participants had to be between 18 and 45 years of age and completed a minimum of 11 years of formal education. Following data screening and cleaning procedures, 35 participants were excluded from the analyses. Specifically, 15 participants did not complete the study, three participants failed to provide any correct responses in the word-fragment completion task, and 17 participants did not follow instructions in the free association task. Among these, six participants failed to provide any responses, while eleven provided invalid responses (e.g., proper nouns, multiple-word answers, or disallowed content such as brand names). The final sample consisted of 85 participants (53 males, 31 females, 1 preferred not to disclose), with a mean age of 24.89 years (*SD* = 5.87; range = 18–43). On average, participants had completed 16.62 years of education (*SD* = 3.02). The majority of participants (75.5%) were students, and most reported no history of drug use or current mental illness diagnosis (see Table 1A in supplementary materials).

**Table 1.** Correlation Coefficients between GTP Frequency Scores and Other Variables.

|  |  | 1 | 2 | 3 | 4 | 5 | 6 | 7 | 8 | 9 | 10 | 11 | 12 | 13 |
|---|---|---|---|---|---|---|---|---|---|---|---|---|---|---|
| 1. | GTP | 1 |  |  |  |  |  |  |  |  |  |  |  |  |
| 2. | WFC Task | .002 | 1 |  |  |  |  |  |  |  |  |  |  |  |
| 3. | AUT | .297** | .114 | 1 |  |  |  |  |  |  |  |  |  |  |
| 4. | FA Task | .095 | .041 | -.029 | 1 |  |  |  |  |  |  |  |  |  |
| 5. | IGD | .552** | .087 | -.02 | .146 | 1 |  |  |  |  |  | \| |  |  |
| 6. | Hours per Week | .217** | -.044 | -.121 | .133 | .305** | 1 |  |  |  |  |  |  |  |
| 7. | Hours per Sess. | .358** | -.078 | -.036 | .091 | .295** | .624** | 1 |  |  |  |  |  |  |
| 8. | Years of Playing | -.051 | .000 | .193 | .136 | -.116 | .065 | -.014 | 1 |  |  |  |  |  |
| 9. | Mind-Popping | .364* | .027 | .416** | .026 | .309* | .020 | .055 | .262* | 1 |  |  |  |  |
| 10. | Daydreaming | .499** | -.048 | .324** | .147 | .397** | .125 | .163 | .182 | .391** | 1 |  |  |  |
| 11. | Pos_Schiz | .607** | -.074 | .171 | .124 | .377**. | -.011 | .161 | -.212 | .392** | .428** | 1 |  |  |
| 12. | Dis_Schiz | .350** | -.012 | .090 | .168 | .437** | .104 | .247* | .009 | .381* | .502** | .383** | 1 |  |
| 13. | Neg_Schiz | .11 | -011 | .-.043 | ,015 | .148 | .013 | .226* | -.066 | -.001 | .108 | .010 | 192 | 1 |

**Note:** GTP: Game Transfer Phenomena (total frequency score); WFC Task: Word Fragment Completion Task (priming score); AUT: Alternate Uses Task; FA Task: Free Association Task (total forward flow coefficient); IGD: Internet Gaming Disorder; Hours per Week: Hours Played per Week; Hours per Sess: Hours Played per Session; Years of Playing: Number of Years of Gaming; Mind-popping frequency; Daydreaming frequency; Pos\_Schiz: Positive Schizotypy; Dis\_Schiz: Disorganised Schizotypy; Neg\_Schiz: Negative Schizotypy
\* p < 0.05, \*\* p < .001

Data collection took place from August 2, 2021, to April 30, 2022. Participants were recruited through official university platforms and social media adverts. Detailed demographic characteristics and gaming-related variables are presented in supplementary materials in Tables 1A and 2A. Overall, participants reported a substantial number of years of gaming experience (*M* = 15.94, *SD* = 7.71), with an average of 18.74 hours of gameplay per week (*SD* = 12.64). Most participants identified themselves as mid-core or hardcore gamers. The study received approval from the University Ethics Committee (protocol number: LMS/PGT/UH/03887). In accordance with the Declaration of Helsinki, all participants provided written informed consent digitally (via online form) prior to taking part in the study. As a token of appreciation for their time and participation, all participants who completed the study were entered into a raffle for a chance to win several £10 and £20 Amazon digital vouchers.

**Table 2.** Hierarchical Multiple Linear Regression for Variables Predicting the Outcome Variable of GTP Frequency (Four Blocks)

|  |  | *Unstandardized coefficients* |  | *Standardized coefficients* |  |  |  |  |  |  |  |
| *Block* | *Predictors* | B | SE | β | *p-value* | R<sup>2</sup> | *Adjusted R<sup>2</sup>* | *ΔR<sup>2</sup>* | *F* | *p-value* | *VIF* |
|---|---|---|---|---|---|---|---|---|---|---|---|
| 1 | Hours per week | -.110 | .116 | -.110 | .343 | .354 | .330 | .354 | 14.80 | .001*** | 1.681 |
|  | Hours per session | 2.128 | .882 | .278 | .018* |  |  |  |  |  | 1.670 |
|  | Internet gaming disorder | .887 | .167 | .504 | .001*** |  |  |  |  |  | 1.125 |
| 2 | Hours per week | -.088 | .108 | -.088 | .420 | .454 | .419 | .100 | 7.22 | .001** | 1.690 |
|  | Hours per session | 1.960 | .823 | .256 | .020* |  |  |  |  |  | 1.677 |
|  | Internet gaming disorder | .620 | .170 | .352 | .001** |  |  |  |  |  | 1.358 |
|  | Mind-popping | .632 | .433 | .135 | .147 |  |  |  |  |  | 1.230 |
|  | Daydreaming | .337 | .117 | .276 | .005* |  |  |  |  |  | 1.317 |
| 3 | Hours per week | -.013 | .101 | -.013 | .897 | .560 | .520 | .106 | 9.28 | .001*** | 1.825 |
|  | Hours per session | 1.577 | .771 | .206 | .044* |  |  |  |  |  | 1.849 |
|  | Internet gaming disorder | .512 | 1.63 | .291 | .002** |  |  |  |  |  | 1.455 |
|  | Mind-popping | .325 | .411 | .069 | .431 |  |  |  |  |  | 1.302 |
|  | Daydreaming | .261 | .115 | .213 | .027* |  |  |  |  |  | 1.416 |
|  | Disorganised schizotypy | -.379 | .332 | -.109 | .258 |  |  |  |  |  | 1.453 |
|  | Positive schizotypy | 1.715 | .405 | .387 | .001*** |  |  |  |  |  | 1.109 |
| 4 | Hours per week | .000 | .099 | .000 | .998 | .582 | .538 | .022 | 4.00 | .049* | 1.833 |
|  | Hours per session | 1.547 | .757 | .202 | .044* |  |  |  |  |  | 1.850 |
|  | Internet gaming disorder | .572 | .163 | .325 | .001*** |  |  |  |  |  | 1.516 |
|  | Mind-popping | .004 | .434 | .001 | .993 |  |  |  |  |  | 1.496 |
|  | Daydreaming | .253 | 1.13 | .207 | .028* |  |  |  |  |  | 1.416 |
|  | Disorganised schizotypy | -.325 | .327 | -.093 | .324 |  |  |  |  |  | 1.540 |
|  | Positive schizotypy | 1.517 | .409 | .342 | .001*** |  |  |  |  |  | 1.111 |
|  | Alternate Uses Task | 1.247 | .623 | .172 | .049* |  |  |  |  |  | 1.343 |

*Note. N= 85; ΔR<sup>2</sup> = R<sup>2</sup> Change, \* = p< .05, \*\* = p<.01, \*\*\* = p<.001*

## Materials and Measures

### Cognitive Tasks

#### Word Fragment Completion Task (WFC)

Implicit memory (priming) was assessed using a word fragment completion task adapted from Roediger [39]. The stimuli were selected from the normative database of 192 single-solution word fragments developed by Erickson et al. [41], which were normed in a sample of 80 college students and controlled for familiarity and baseline completion probability. Two equivalent word lists (Lists A and B) were created and counterbalanced across participants (see Supplementary Materials for the complete word lists). Each list comprised 20 words and was matched on word length (List A: *M* = 7.45, *SD* = 0.61; List B: *M* = 7.45, *SD* = 0.69), familiarity (List A: *M* = 6.20, *SD* = 0.90; List B: *M* = 6.37, *SD* = 1.10), and baseline fragment completion probability (List A: *M* = 0.13, *SD* = 0.10; List B: *M* = 0.17, *SD* = 0.14).

In the study phase, half of the participants rated each word from List A and another half from List B for pleasantness using a 5-point Likert scale (1 = *very unpleasant*, 2 = *unpleasant*, 3 = *neither pleasant nor unpleasant*, 4 = *pleasant*, 5 = *very pleasant*). Each trial was preceded by a 1.5 sec fixation point in the centre of a screen followed by a word presented on the screen for 6 sec together with the rating scale for pleasantness rating. During the test phase, all participants completed 40 word fragments created from words in List A and List B. Thus, for each participant, 20 fragments corresponded to previously studied words and 20 fragments to novel (unstudied) words: ellipse ( _ l l _ p _ e), gazzelle (_ a z _ l _ e), pendulum (_ e _ d _ l _ m), dinosaur (d _ n _ s a _ _). Each fragment was presented for 12 seconds, during which participants were required to provide the first solution that came to mind (see Figure 1 for a timeline of study and test phase). Priming performance was operationalized as the difference between the proportion of correctly completed studied fragments and unstudied fragments, with higher scores indicating stronger implicit memory effects, and the score of 0 representing no priming.

**Figure 1.**
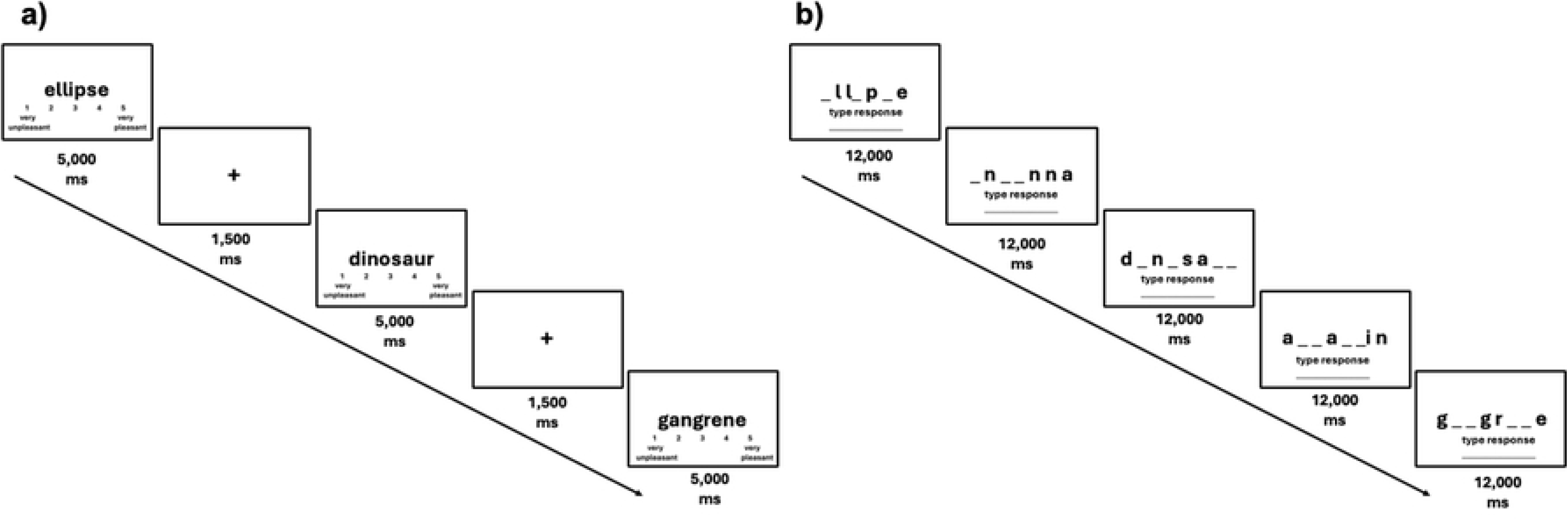
Word Fragment Completion Task: Study (Panel a) and Test (Panel b) phase timeline

#### Alternate Uses Task (AUT)

Creativity was assessed using the Alternate Uses Task [31], with the objects “car tyre” and “newspaper” serving as seed items. These objects were selected due to their frequent use in creativity research and their suitability for generating diverse responses.

Participants were instructed to generate as many unusual and creative uses for each object as possible within a 2-minute time limit per item. To encourage originality, participants were explicitly instructed to avoid typical or commonly known uses, consistent with previous findings that individuals tend to rely on easily accessible and familiar information during idea generation [34]. Common examples for ‘car tyre’ would be flowerpot, swing, racetrack, and for “newspaper” – wrapping items, covering surfaces, starting a fire, etc.

Furthermore, responses were scored using a uniqueness index based on the method proposed by Wallach and Kogan [32]. Each response received 1 point if it was unique within the sample (i.e., provided by only one participant) and 0 points if it was repeated by others. To improve scoring validity, an additional usability criterion was applied [33], whereby responses that were unique but impractical or implausible were assigned 0 points. The total creativity score was computed by summing all valid unique responses across both items.

#### Free Association Task

Associative thinking and divergent thinking were further assessed using a free association task based on the forward flow paradigm [35]. Participants were presented with two initial cue words (“bear” and “table”). For each cue word, they were instructed to generate a chain of 19 consecutive single-word associations. Each response served as the cue for the subsequent response. Participants were instructed to respond spontaneously, without deliberate effort to control or optimize their responses, and to avoid the use of proper nouns or multi-word expressions. No time limit was imposed. Divergent thinking was quantified using the forward flow metric, which measures the average semantic distance between each word and all preceding words in the sequence using latent semantic analysis. Scores range from 0 (minimal semantic distance) to 1 (maximum semantic distance), with higher values reflecting greater divergence in associative thinking. Forward flow scores were computed using the open-access platform provided by Gray et al. [35].

### Questionnaires

#### Gaming habits questionnaire

Gaming habits were assessed using a 15-item questionnaire that measured gaming experience (e.g., years of playing), weekly gaming time, typical session duration, gaming proficiency, etc.

#### Game Transfer Phenomena Scale (GTPS)

The 20-item Game Transfer Phenomena Scale [42] measures the frequency of GTP experiences during the previous 12 months. The scale comprises five dimensions: altered visual perceptions, altered body perceptions, altered auditory perceptions, automatic mental processes, and behaviours and actions, with four items per dimension. The scale demonstrated excellent internal consistency (Cronbach’s α = .94) as well as cross-validation and criterion-related validity [42]. Responses were recorded on a 5-point Likert scale (0 = *Never*, 1 = *Once or twice*, 2 = *A few times*, 3 = *Several times*, 4 = *Many times*) [6].

#### Daydreaming Frequency Scale (DDFS)

Daydreaming frequency was measured using the 12-item Daydreaming Frequency Scale (DDFS), a subscale of the Imaginal Processes Inventory [43,44]. The scale assesses the frequency of spontaneous task-unrelated thoughts and mind-wandering in everyday life. Items are rated on a 5-point Likert scale, with total scores ranging from 12 to 60, where higher scores indicate more frequent daydreaming.

#### Mind-Popping Questionnaire (MPQ-4)

The Mind-Popping Questionnaire [8] was used to assess the occurrence and frequency of involuntary semantic memories (mind-pops). These refer to fragments of general knowledge such as a word, saying, image or music popping into mind, often without any obvious internal of external triggers. The questionnaire first determines whether participants have ever experienced mind-popping and, if so, assesses its frequency using an 8-point response scale ranging from 1 = *only a few times in my entire life* to 8 = *three or more times a day*.

Additional items assess the types of mind-pops experienced and allow participants to describe examples in their own words. In the present study, only the frequency measure was analysed, yielding scores ranging from 0 (participants reporting no experience of mind-popping) to 8 (participants reporting occurrences three or more times per day).

#### Internet Gaming Disorder Scale–Short Form (IGDS9-SF)

Symptoms of Internet Gaming Disorder were assessed using the 9-item Internet Gaming Disorder Scale–Short Form [45], which evaluates the nine diagnostic criteria proposed by the American Psychiatric Association [46] . Responses are provided on a 5-point Likert scale (ranging from 1 = *Never* to 5 = *Very often*), producing total scores between 9 and 45, with higher scores indicating greater severity of gaming disorder symptoms. The instrument has demonstrated good psychometric properties and cross-cultural validity.

#### Multidimensional Schizotypy Scale–Brief (MSS-B)

Schizotypal traits were assessed using the 38-item Multidimensional Schizotypy Scale–Brief [47], which uses a dichotomous (Yes/No) response format. The scale measures three dimensions of schizotypy: positive (13 items), negative (13 items), and disorganized (12 items). Total scores were computed separately for each dimension after reverse-scoring items 4, 10, 25, and 37, with higher scores indicating greater levels of schizotypal traits. The MSS-B is a brief version of the original Multidimensional Schizotypy Scale and has demonstrated good cross-validation and psychometric properties.

#### Procedure

The study questionnaires and cognitive tasks were presented online using the Qualtrics software. Participants who expressed interest in taking part, received a digital information sheet outlining the study’s purpose and procedures. After providing informed consent electronically, participants were assigned a unique anonymous identifier and given remote access to the study via a personalized link.

All participants completed the tasks in a fixed order. First, participants completed the demographic and gaming habits questionnaires. This was followed by the free association task, designed to capture spontaneous thought processes early in the session. Participants then completed the GTP scale, followed by a word-fragment completion task comprising two phases; the encoding (study) phase where participants were instructed to rate the pleasantness of 20 words, each presented for a fixed duration of 6 seconds, irrespective of participants’ response latency, to ensure that all participants were exposed to primes for the same amount of time. During the delay phase (approximately 25–30 minutes), participants completed the Alternate Uses Task and a series of questionnaires assessing schizotypy, IGD, mind-popping, and daydreaming frequency. Finally, participants completed the test phase of the word fragment completion task. During this phase, participants completed a word fragment completion task consisting of 46 items (6 fragments were used for practice). Instructions explicitly stated that they would be presented with word fragments one by one and were required to type the correct word to complete the blanks. All target words were restricted to singular nouns, excluding proper names or brands. Each trial had a time limit of 12 seconds for completion. Participants were instructed to try their best as some fragments were challenging; if they completed a fragment before the time expired, they were instructed to click an arrow button to proceed immediately to the next word fragment. The entire experimental session lasted approximately 45 to 60 minutes. Upon completion, participants were debriefed and compensated with an entry into a raffle for monetary vouchers.

## Results

### Data Analysis

Data were screened and cleaned for completeness and adherence to task instructions prior to analysis. Pearson correlations were conducted to examine associations between priming scores, creativity measures, GTP, and related constructs. Multiple regression analyses were performed to assess the predictive value of creativity, divergent thinking, schizotypy, and IGD on GTP frequency.

### Priming Effects

Priming scores were computed by subtracting the proportion of correctly completed fragments for non-studied words from the proportion of correctly completed fragments for previously studied words. Higher scores therefore reflect stronger implicit priming effects. The majority of participants (91.7%, *n* = 78) demonstrated a positive priming effect. The mean priming score was *M* = .225 (*SD* = .142), with values ranging from −.05 to .50. Only five participants showed no priming effect, and two participants showed a small negative effect (−.05). Overall, the distribution of priming scores indicated a robust implicit memory effect across the sample (see Figure 2).

**Figure 2.**
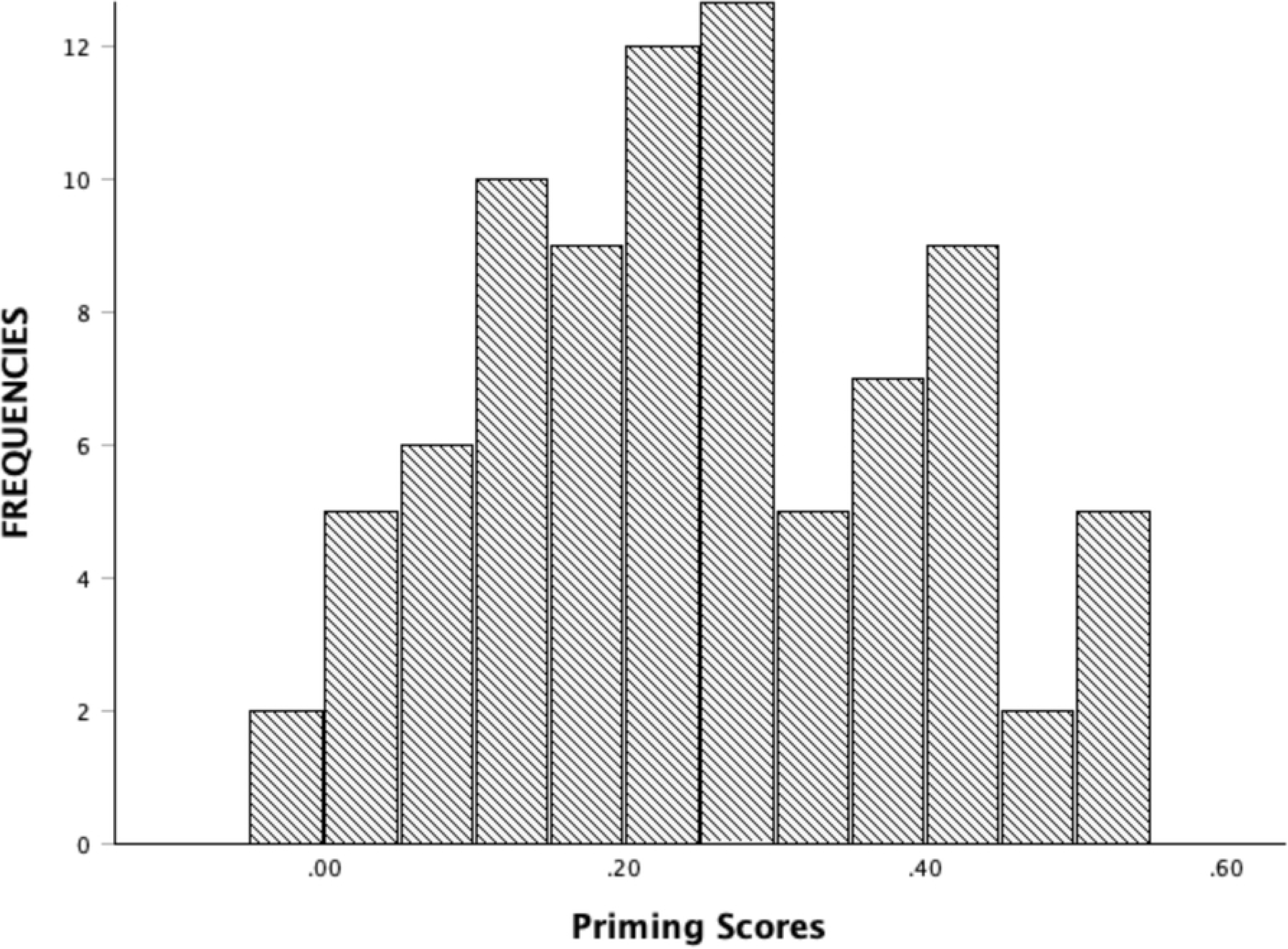
Frequency Distribution of Participant Priming Scores

### Alternate Uses Task (AUT)

Participants generated a total of 776 responses across the two AUT cue objects, with 362 responses for ‘car tyre’ and 415 for ‘newspaper’. After applying the criteria of uniqueness and usability, 158 responses were retained as valid creative responses. For ‘car tyre’ participants produced an average of 0.74 (*SD* = 0.92) unique and useful responses (range = 0– 4), whereas for ‘newspaper’, the mean was 1.11 (*SD* = 1.15; range = 0–4). The combined mean AUT score, calculated as the total number of unique and useful responses across both items, was 1.77 (*SD* = 1.73), with a range of 0 to 8. This composite score was used in all subsequent analyses.

### Free association Task

Participants showed relatively high levels of divergent thinking calculated through the forward flow coefficient in the free association task. For the cue word ‘bear’, the mean was .78 (*SD* = .058; range = .60–.90), and for ‘table’, the mean was .80 (*SD* = .053; range = .64–.90). These results suggest a generally high degree of semantic divergence in associative thinking across participants. Figure 3 illustrates representative examples of high and low forward flow sequences within the sample.

**Figure 3.**
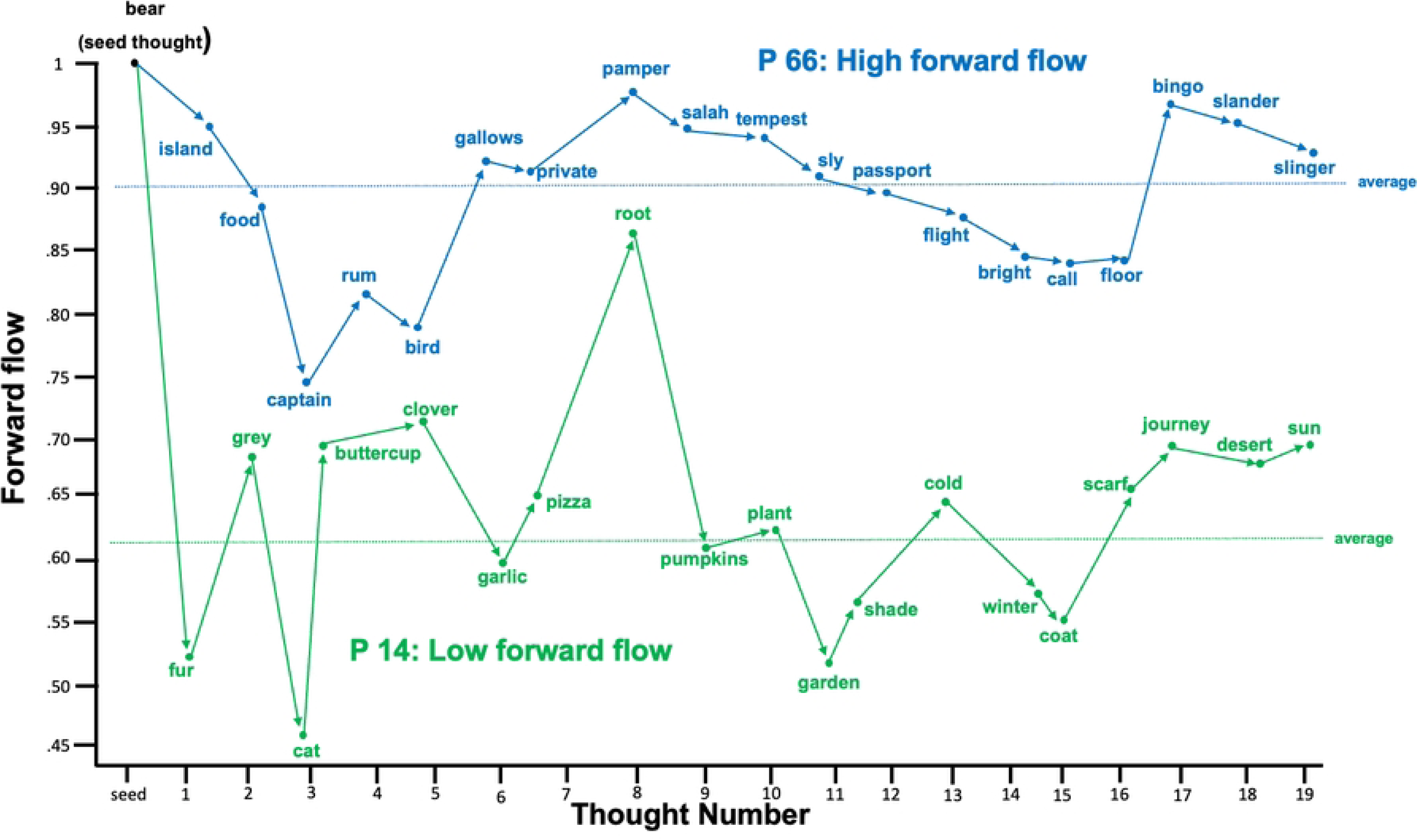
Thought plot comparing two participants: responses from Participant 66 (blue) indicate a high forward flow coefficient, whereas responses from Participant 14 (green) reflect a low forward flow coefficient.

### Correlational Analyses

Pearson bivariate correlations were conducted to examine associations between GTP frequency and all independent variables, including gaming behaviour, spontaneous cognitive processes, schizotypal traits, and cognitive task performance (see Table 1).

GTP frequency was positively correlated with AUT scores, internet gaming disorder (IGD), hours played per session, mind-popping frequency, daydreaming frequency, positive schizotypy, and disorganized schizotypy (all *p* < .01), as well as with hours played per week (*p* < .05). No significant associations were found between GTP and WFCT (priming scores), forward flow coefficient, years of playing, or negative schizotypy. Regarding relationships among independent variables, mind-popping frequency was positively correlated with AUT scores, daydreaming frequency, positive schizotypy, and disorganized schizotypy.

Daydreaming frequency was also positively associated with positive and disorganized schizotypy, but not with the creativity measures (i.e., AUT scores or forward flow coefficient).

### Results of hierarchical regression analysis (4 Blocks)

Prior to analysis, the assumptions of regression were evaluated, and no violations were detected; multicollinearity was low overall. To examine the direct associations between GTP frequency and gaming-related, psychopathological, and cognitive variables, a four-block hierarchical multiple regression was conducted with total GTP score as the dependent variable. Gaming variables (hours played per week, hours per session, and IGD) were entered in Block 1; mind-popping and daydreaming in Block 2; disorganized and positive schizotypal traits in Block 3; and AUT total score (creativity) in Block 4.

In Block 1, gaming variables accounted for 35.4% of the variance in GTP (p < .001), with hours per session and IGD emerging as significant predictors. The inclusion of mind-popping and daydreaming in Block 2 explained an additional 10% of variance (R² = .454, ΔR² = .100, p < .001), with hours per session, IGD, and daydreaming remaining significant. In Block 3, the addition of disorganized and positive schizotypal traits accounted for a further 10.6% of variance (R² = .560, ΔR² = .106, p < .001), with hours per session, IGD, daydreaming, and positive schizotypy as significant predictors. Finally, in Block 4, creativity (AUT) contributed an additional 2.1% of variance (R² = .582, ΔR² = .022, p < .049); in this final model, hours per session, IGD, daydreaming, positive schizotypy, and AUT scores were significant predictors of GTP.

Overall, the full model explained 58.2% of the variance in GTP frequency. Consistent with previous studies IGD, daydreaming frequency, and positive schizotypy emerged as the strongest predictors. Additionally, hours played per session and AUT performance were identified as new significant predictors in the present study. Detailed results for each step, including standardized beta coefficients and significance levels, are reported in Table 2.

To determine whether specific GTP dimensions (Altered Perceptions, Automatic Mental Processes, and Behaviours and Actions) drove these effects, separate linear regressions were conducted for each dimension as an outcome variable. All models were significant (p < .001).

## Discussion

The present study investigated, for the first time, the relationship between implicit memory (priming), creativity, and Game Transfer Phenomena (GTP), while also examining previously identified predictors of GTP, including Internet Gaming Disorder (IGD), spontaneous cognitive processes, and schizotypal traits. Several key findings emerged. First, although participants demonstrated robust perceptual priming effects at the group level, individual differences in priming performance were not associated with self-reported GTP frequency. Second, creativity, as measured by the Alternate Uses Task (AUT), was positively associated with GTP and contributed uniquely to its prediction, whereas the forward flow coefficient showed no significant relationships. Third, consistent with previous studies, IGD, positive schizotypy, and daydreaming emerged as the strongest predictors of GTP, alongside hours played per session.

### Absence of association between priming and GTP

Despite clear evidence of perceptual priming at the group level, individual differences in priming performance were unrelated to GTP frequency. This finding does not support the hypothesis that perceptual domain-general implicit memory sensitivity underlies susceptibility to GTP experiences. At first glance, this finding appears inconsistent with previous research on involuntary autobiographical memories and mind-popping, where priming mechanisms have been proposed to facilitate the spontaneous activation of previously encoded information [8,21]. Moreover, a recent diary-based study on GTP has shown that these experiences tend to occur shortly after gameplay (often within 24 hours) and are frequently triggered by identifiable internal or external cues that are identical to gaming stimuli or conceptually related to them, providing converging evidence that recent gameplay can produce relatively long-lasting activation effects [7]. Thus, the present null findings should not be interpreted as evidence against an activation-based account of GTP, but rather as suggesting that the specific form of implicit memory measured may be critical.

For example, it may be important to consider the type of priming that is being assessed. The word-fragment completion task, employed in the present study, primarily measures perceptual and lexical implicit memory processes involving relatively constrained repetition-based facilitation effects. In contrast, GTP could depend more strongly on broader semantic and autobiographical activation mechanisms derived from prolonged and immersive gameplay exposure. This interpretation is consistent with contemporary semantic-to-autobiographical priming framework [24,27], which proposes that spontaneous cognitions emerge through spreading activation across interconnected semantic and autobiographical memory networks. Supporting this view, Mace and Aaron [27] demonstrated that narrative processing itself can increase the subsequent occurrence of involuntary autobiographical memories, suggesting that autobiographical activation may emerge from meaningful semantic contexts rather than from simple perceptual repetition alone.

Similarly, Coane and Balota [28] showed that repeated naturalistic exposure to highly interconnected lexical concepts (e.g., holidays and festivities) can maintain elevated accessibility over extended periods of time outside laboratory settings. This may be particularly relevant for GTP, as video games involve repeated multisensory exposure, emotional salience, narrative engagement, and highly immersive cognitive processing. Consequently, GTP may emerge less from individual differences in basic perceptual priming ability and more from sustained activation of gaming-related semantic and autobiographical representations embedded within everyday cognition.

From this perspective, the absence of associations between word-fragment performance and GTP frequency may reflect a mismatch between the cognitive processes measured by the task and the mechanisms actually involved in GTP experiences. Whereas the present word-fragment completion task captured relatively low-level perceptual implicit memory processes, GTP may rely more strongly on conceptual semantic and autobiographical activation processes, which could be assessed by conceptual priming tasks (e.g., a category exemplar task, etc.). Such tasks may provide a more direct test of whether GTP are associated with semantic-to-autobiographical priming mechanisms.

### Creativity and GTP

Creativity, as measured by the Alternate Uses Task (AUT), was positively associated with GTP frequency and remained a significant predictor even after controlling for gaming behaviour and psychopathological traits. This finding is consistent with previous research suggesting that individuals prone to fantasy and imagination tend to exhibit higher levels of creativity [48] and with studies linking GTP-related experiences to fantasy proneness [49,50].

One possible interpretation is that creativity reflects a broader cognitive style characterized by flexible associative thinking and reduced constraints on semantic activation, which may increase access to remote or weakly connected semantic representations, thereby facilitating unusual cognitive associations and increasing susceptibility to GTP experiences [51,52]. Contemporary semantic network accounts of creativity propose that highly creative individuals navigate semantic memory more flexibly, allowing activation to spread more readily between distant concepts and increasing the likelihood of forming novel associations [52]. This interpretation is further supported by the observed associations between AUT scores, mind-popping, and positive schizotypy, consistent with previous findings linking involuntary semantic memories and creative cognition [30]. Similar associations between involuntary semantic cognitions and psychosis-proneness have been reported in schizophrenia research, where increased frequencies of mind-pops have been observed among individuals with schizophrenia [53,54].

These findings align with theoretical accounts proposing that creativity and schizotypy partially overlap through mechanisms involving increased access to remote or loosely connected semantic representations [36]. However, the relationship between creativity and schizotypy remains complex and inconsistent across studies, likely due to differences in operationalization and the multidimensional nature of both constructs [36]. Importantly, no association was found between creativity and daydreaming, which is consistent with mixed findings in the literature on mind-wandering and creativity [11]. This suggests that not all forms of spontaneous cognition are equally related to creative thinking.

Contrary to expectations, the forward flow coefficient did not correlate with creativity, GTP, or spontaneous cognitive variables. This finding was unexpected given previous evidence linking forward flow to divergent thinking and creative cognition [35]. Several methodological factors may explain this null finding. First, the study was conducted entirely online, reducing experimental control over participants’ testing environments and potentially increasing distractions during task completion. Second, free association paradigms may be particularly sensitive to contextual noise and fluctuations in participant engagement compared to more structured cognitive tasks. Unlike structured divergent thinking tasks, free association tasks paradigms rely heavily on spontaneous responding and sustained adherence to instructions, which may be more difficult to ensure in unsupervised settings. Future studies should therefore examine forward flow under more controlled laboratory conditions to better evaluate its relationship with GTP and spontaneous cognition.

### Predictors of GTP

The hierarchical regression analyses revealed a consistent pattern of predictors. Across all models, IGD, positive schizotypy, and daydreaming frequency emerged as the strongest predictors of GTP, replicating findings from previous studies [6,7].

Additionally, hours played per session emerged as a significant predictor in the present study. This finding suggests that the intensity and duration of individual gameplay sessions, rather than cumulative gaming experience alone, may be particularly relevant for the occurrence of GTP. This interpretation aligns with previous research linking prolonged gaming exposure to cognitive and perceptual aftereffects [50].

Importantly, the role of daydreaming further supports the conceptualization of GTP as part of a broader class of spontaneous cognitive phenomena characterized by involuntary and internally generated experiences. Moreover, the associations observed between mind-popping, daydreaming, and positive schizotypy suggest the presence of shared mechanisms involving increased cognitive permeability and reduced filtering of internally generated content. Taken together, these findings further suggest that GTP are unlikely to be explained by a single cognitive mechanism. Rather, they appear to emerge from the interaction between prolonged gaming exposure, semantic-autobiographical memory activation, individual differences in cognitive style, and vulnerability to spontaneous internally generated cognition.

### Limitations and Future Research

Several limitations should be considered when interpreting the present findings. First, an important limitation concerns the operationalization of priming. The word-fragment completion task employed in the present study primarily indexes perceptual and lexical implicit memory processes and is widely used as a measure of perceptual priming. However, the reliability of implicit memory tasks used to assess individual differences has been found to be lower than that of explicit memory measures, potentially reducing the magnitude of associations with other constructs [55]. Moreover, evidence from previous research on GTP suggests that these experiences may be more closely related to conceptual or semantic forms of long-term priming than to perceptual repetition effects. Consequently, the absence of associations between perceptual priming scores and GTP frequency should not be interpreted as evidence against the involvement of priming mechanisms in GTP more generally. Rather, the present findings suggest that perceptual priming, as assessed here, may not represent the form of priming most relevant to these experiences. Future studies should therefore employ conceptual and semantic priming paradigms to further clarify the role of long-term priming processes in GTP.

Second, data collection was conducted entirely online due to COVID-19 restrictions, reducing experimental control over participants’ testing environments and levels of engagement. Nevertheless, online methodologies have become well established in cognitive and individual differences research and enabled the recruitment of a larger and more diverse community samples during a period in which laboratory testing was not feasible. Although uncontrolled distractions may have introduced additional variability into task performance, such influences would be expected to attenuate rather than artificially inflate observed associations. Therefore, the significant associations observed in the present study are unlikely to be solely attributable to characteristics of the online testing context.

Finally, the cross-sectional design and reliance on retrospective self-report measures limit causal inferences and may be susceptible to memory biases. Consequently, it remains unclear whether intensive gaming contributes to the development of cognitive-perceptual tendencies associated with GTP, or whether individuals with a pre-existing susceptibility to spontaneous and unusual mental experiences are more likely to report both problematic gaming behaviours and GTP. Longitudinal and experimental studies employing multiple methods of assessment will be necessary to clarify the directionality of these relationships and further elucidate the cognitive mechanisms underlying GTP.

## Conclusions

The present study provides new insights into the cognitive and psychological mechanisms underlying GTP. Although individual differences in perceptual priming scores were not associated with GTP frequency, creativity as assessed by participants AUT scores emerged as a novel correlate of GTP. In addition, the consistent finding that IGD, positive schizotypy, and daydreaming were key predictors of GTP frequency reinforces the conceptualization of GTP as arising from the interaction between gaming behaviour, personality traits, and spontaneous mental processes.

Overall, these findings contribute to a more integrated understanding of GTP and highlight the importance of considering both cognitive mechanisms and individual differences in explaining the phenomenon. Future research should employ more ecologically valid and experimentally controlled paradigms to further clarify the mechanisms underlying GTP experiences.

## Acknowledgements

The research, presented in this study, was supported by a PhD studentship awarded to the first author by the Mexican National Council for Humanities, Science and Technology (award number 709988) to conduct research on GTP. The authors are also grateful for the support provided by the School of Life and Medical Sciences at the University of Hertfordshire and to all the video game players who participated in the study.

## Appendix

*Examples of Game Transfer Phenomena (GTP) Experiences Reported by Players*

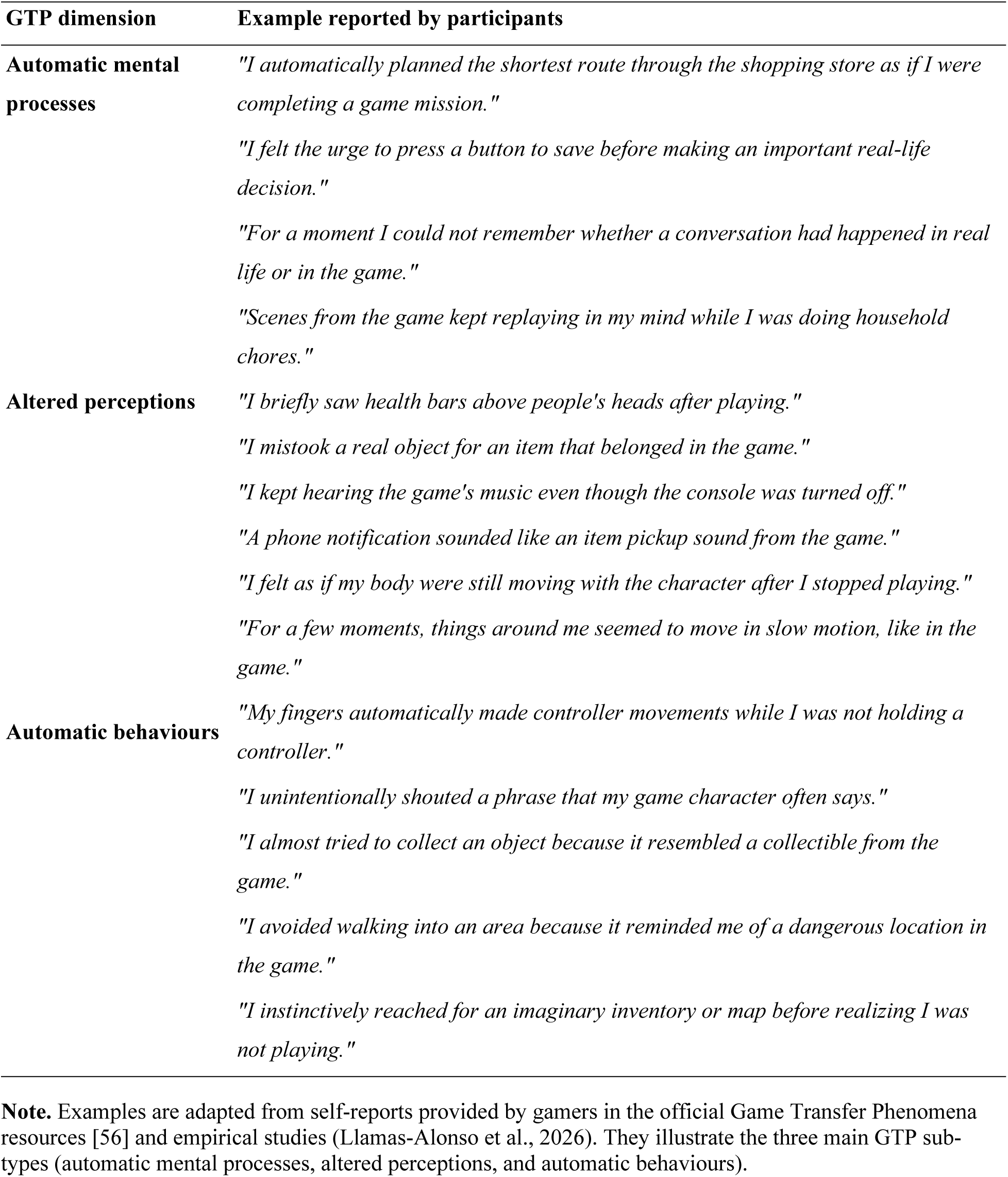

| **GTP dimension** | **Example reported by participants** |
|---|---|
| **Automatic mental processes** | *"I automatically planned the shortest route through the shopping store as if I were completing a game mission."* |
|  | *"I felt the urge to press a button to save before making an important real-life decision."* |
|  | *"For a moment I could not remember whether a conversation had happened in real life or in the game."* |
|  | *"Scenes from the game kept replaying in my mind while I was doing household chores."* |
| **Altered perceptions** | *"I briefly saw health bars above people's heads after playing."* |
|  | *"I mistook a real object for an item that belonged in the game."* |
|  | *"I kept hearing the game's music even though the console was turned off."* |
|  | *"A phone notification sounded like an item pickup sound from the game."* |
|  | *"I felt as if my body were still moving with the character after I stopped playing."* |
|  | *"For a few moments, things around me seemed to move in slow motion, like in the game."* |
| **Automatic behaviours** | *"My fingers automatically made controller movements while I was not holding a controller."* |
|  | *"I unintentionally shouted a phrase that my game character often says."* |
|  | *"I almost tried to collect an object because it resembled a collectible from the game."* |
|  | *"I avoided walking into an area because it reminded me of a dangerous location in the game."* |
|  | *"I instinctively reached for an imaginary inventory or map before realizing I was not playing."* |

**Note.** Examples are adapted from self-reports provided by gamers in the official Game Transfer Phenomena resources [56] and empirical studies (Llamas-Alonso et al., 2026). They illustrate the three main GTP sub-types (automatic mental processes, altered perceptions, and automatic behaviours).

